# BatchSVG: identifying batch-biased genes in the application of spatially variable gene detection

**DOI:** 10.64898/2025.12.09.693192

**Authors:** Kinnary Shah, Christine Hou, Jacqueline R. Thompson, Stephanie C. Hicks

## Abstract

A standard task in the analysis of spatially resolved transcriptomics data is to identify spatially variable genes (SVGs). This is most commonly done within one tissue section at a time because the spatial relationships between the tissue sections are typically unknown. However, large-scale spatial atlases are being generated, for example across hundreds of donors, where the goal is to identify a common set of SVGs to use for downstream analyses. One challenge is how to identify and remove SVGs that are associated with a known bias or technical artifact, such as the slide or capture area, which can lead to poor performance in downstream analyses, such as spatial domain detection. Here, we introduce `BatchSVG`, a tool to identify batch-biased genes in the application of SVG detection. Our approach compares the rank of per-gene deviance under a binomial model (i) with and (ii) without including a covariate in the model that is associated with the known bias or technical artifact. If the rank of a gene changes significantly between these, then we infer that this gene is likely associated with the bias or technical artifact and should be removed from the downstream analysis. We consider two SRT datasets and show how our model can improve the results of downstream analysis.

## 2 Introduction

Recent advances have led to the development of spatially-resolved transcriptomics (SRT) technologies where genes can be measured in a 2D spatial context from individual tissue sections [1]. A common data analysis task with these data is identifying spatial gene expression patterns across a tissue section [2]. These spatially variable genes (SVGs) can be used as input to downstream analyses, such as unsupervised clustering algorithms to identify cell types or spatial domains [3] and enhance the identification of spatial domains over non-spatial feature selection [4, 5].

However, current SVG detection methods do not accommodate multi-sample datasets because there is often no formal spatial relationship between tissue sections. Unlike non-spatial feature selection methods that can model the batch effects that are often present in larger experiments with complex study designs (e.g., donor identity, slide, sequencing batch), it can be challenging to identify SVGs that are associated with unwanted variation or technical artifacts [6].

To address this challenge, we developed `BatchSVG`, a method to identify SVGs that are associated with unwanted technical variation. We demonstrate that batch-biased SVGs can confound clustering results in small, multi-sample experiments. `BatchSVG` builds upon the binomial deviance model [7] and applies a data-driven thresholding approach to refine SVG features used as input to downstream analyses. We implemented `BatchSVG` as an R/Bioconductor package to facilitate broad accessibility and seamless integration into existing SRT analysis pipelines and to ensure compatibility with workflows based on the `SpatialExperiment` framework [8].

## 3 Methods

### 3.1 Overview of BatchSVG methodology

For a given set of tissue sections *t* ∈ (1, …, *T*), we calculate the per-gene residual deviance under a binomial model [7]. We consider all spatial coordinates across tissue sections as independent observations. These models are fit per-gene and the residual deviance is calculated for gene *i* ∈ (1, …, *n*). Generally, a higher per-gene deviance *d*_*i*_ suggests that the gene’s expression is more likely to be biologically meaningful.

To assess the impact of the unwanted variation or batch effect in SRT data, we fit a binomial model per gene (i) with and (ii) without including a covariate in the model that is associated with the batch effect or unwanted variation. Therefore, a reduction in deviance after accounting for the batch covariate indicates that the batch explains a portion of the variation in gene expression that was previously attributed to biological differences. Using this model output above, we define *d*_*i*, default_ and *d*_*i*, batch name_ as the residual deviance for gene *i* using a binomial model with and without the batch effect, respectively. Finally, we calculate a per-gene relative change in deviance (RCD) as 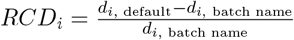

In addition to the residual deviance itself, we also consider the ranks of the residual deviances where top-ranked genes have the largest residual deviance and are considered more important. Here, an increase in rank (e.g. from a rank of 1 to a rank of 500) when including the batch variable indicates that the relative importance of the feature is diminished once the batch variable is accounted for. Therefore, in addition to RCD, we also evaluated the rank deviance (RD), which is defined as *RD*_*i*_ = *r*_*i*, batch name_ − *r*_*i*, default_, where *r*_*i*, default_ and *r*_*i*, batch name_ are the per-gene rank when the binomial deviance model is run with and without the batch variable, respectively.

The continuous nature of RCD and RD complicates the determination of a specific threshold for identifying potentially batch-biased genes. To address this, we developed a data-driven thresholding approach based on the number of standard deviations (nSD) of RCD and RD. The nSD for per-gene RCD is defined as 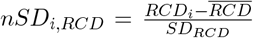, where 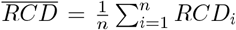 and 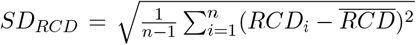. The nSD for per-gene RD is computed using the same formula defined as 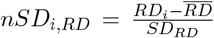, where 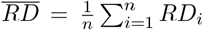 and 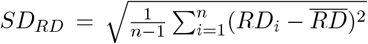. We leverage this standardized measure to establish an adaptive threshold for dataset-specific variability in RCD and RD. This facilitates the identification of genes exhibiting significant deviations due to batch effects.

### 3.2 Evaluation of BatchSVG’s downstream impacts

To evaluate the impact of BatchSVG, we applied `PRECAST` [3] for clustering based on the original feature set and the refined feature set after removing batch-biased SVGs. Clusters were evaluated for consistency with known biology based on known marker gene expression and consistency between individual samples based on the percent abundance. Cluster agreement and similarity was assessed with normalized mutual information (NMI) and principal components analysis used to generate silhouette scores. Common quality control (QC) metrics were evaluated for each cluster from the original features set and the refined features set to examine if differences in clustering outcomes may be addressed by stricter upstream QC filters.

### 3.3 BatchSVG R/Bioconductor package

We implemented the BatchSVG method into the `BatchSVG` R/Bioconductor package, which provides a comprehensive framework to efficiently identify the SVGs impacted by batch effects, such as sample, slide, and sex. The package is designed to work with `SpatialExperiment` objects [8]. The workflow (**Supplementary Figure 1**) includes feature selection (`featureSelect()`), visualization (`svg nSD()`), and finally extraction of batch-biased features (`biasDetect()`). For more details on the functions, please see the vignette on the package website (https://bioconductor.org/packages/BatchSVG).

**Figure 1:**
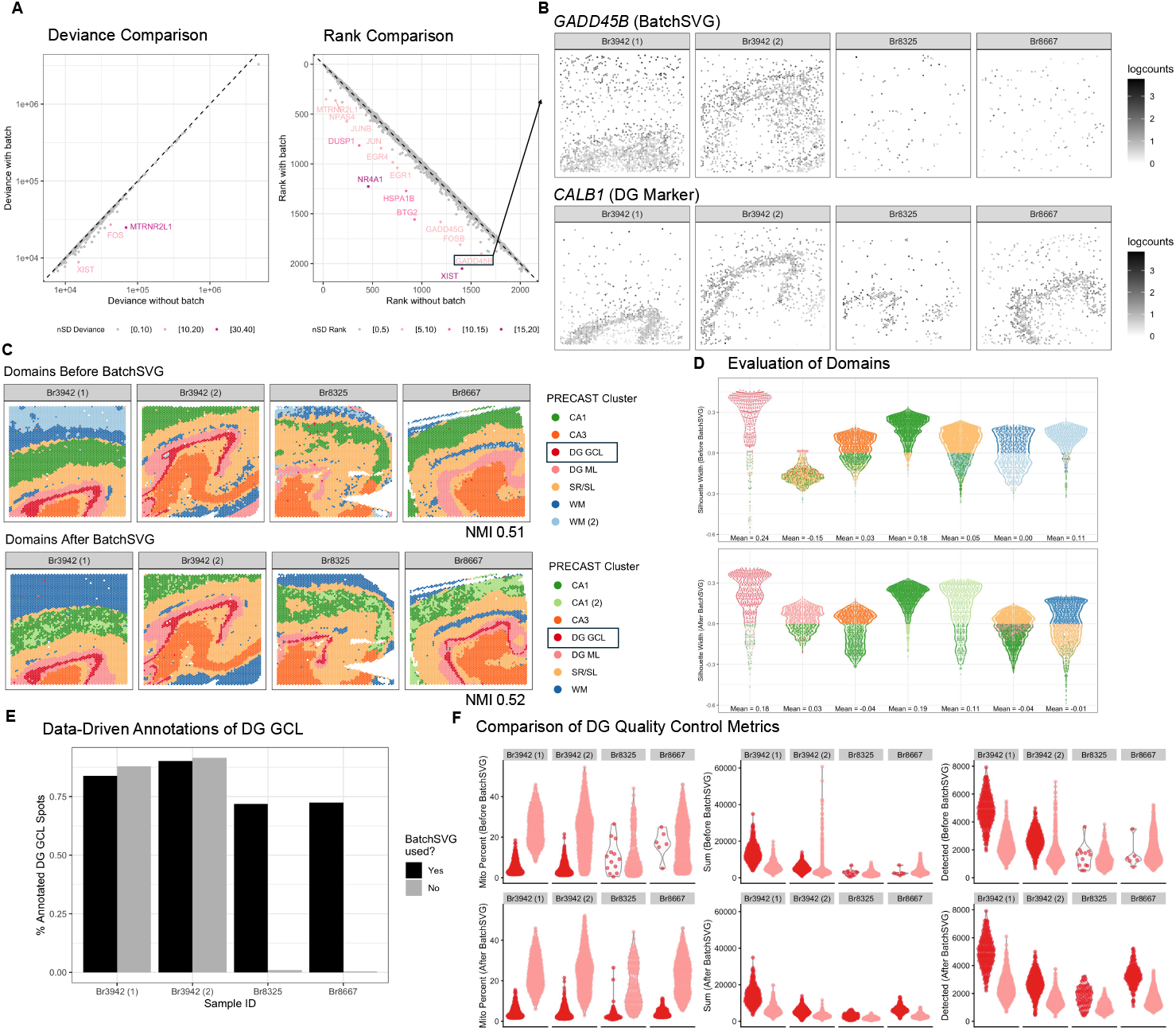
Overview of BatchSVG. (**A**) Left scatterplot displays the change in deviance for each gene. The *x*-axis is deviance calculated without using any covariates. The *y*-axis is deviance calculated with sample as the covariate. The color represents the number of standard deviations of relative change in deviance. Right scatterplot is similar, substituting rank for deviance. (**B**) Top spot plot shows the log-normalized counts gene expression for *GADD45B*, which is a batch-biased SVG boxed in (A). Bottom spot plot shows the log-normalized counts gene expression for *CALB1*, a dentate gyrus (DG) marker. **(C)**Top spot plot shows the clusters derived from `PRECAST` when all SVGs were used as input. Bottom spot plot shows the clusters derived from `PRECAST` when the batch-biased SVGs were excluded from the input SVG list. The clusters were annotated with the following labels - CA: cornu ammonis, DG GCL: dentate gyrus granule cell layer, DG ML: dentate gyrus molecular layer, SR/SL: stratum lucidum and stratum radiatum, WM: white matter. Color represents the cluster. Text displays the normalized mutual information (NMI) for each set of domains compared to previous annotations from the original manuscript. **(D)**Top violin plots display silhouette scores for domains derived from `PRECAST` when all SVGs were used as input. Bottom violin plots display silhouette scores for domains derived from `PRECAST` when the batch-biased SVGs were excluded from the input SVG list. Color represents the closest domain. (**E**) Bar plot displays the percentage of DG GCL spots from the previous annotations that were called as DG GCL spots in either set of `PRECAST` domains. Color represents domains derived from `PRECAST` when the batch-biased SVGs were included or excluded from the input SVG list. (**F**) Violin plots display various quality control metrics for domains derived from `PRECAST` when the batch-biased SVGs were included (top row) or excluded (bottom row) from the input SVG list. Left row displays mitochondrial percent, middle row displays total number of UMIs, and right row displays number of detected genes. Color represents the domain.

## 4 Results

### 4.1 Dentate gyrus identification in human hippocampus

This dataset was generated from adjacent tissue sections of the anterior human hippocampus across ten adult neurotypical donors, using 10x Genomics Visium (n=36 capture areas) [9]. It comprises nine slides with four samples per slide, of which we selected a subset of four samples from the raw SRT dataset. We leveraged the `nnSVG`-based SVG selection process from the original manuscript [9, 10] to include only the top 2,000 ranked features and only genes that appear in more than one sample. With this approach, we obtained 2,082 SVGs across the four samples. Using `BatchSVG` with sample as the batch, we identified 15 batch-biased SVGs (**Figure 1A, Supplementary Table 1**). The spatial expression plots confirmed that these identified features are spatially variable and batch-biased SVGs (**Figure 1B, Supplementary Figure 2**). We defined the refined features set as the 2,082 original SVGs minus the 15 batch-biased SVGs to get 2,067 SVGs. Utilizing *k* = 7 clusters, we found that clusters based on the original feature set and the refined feature set described known spatial domains in the human hippocampus (**Figure 1C, Supplementary Figures 3 and 4**). We highlight the red dentate gyrus granule cell layer (DG GCL) domains, which are outlined by the *CALB1* marker gene (**Figure 1B**). Although application of `BatchSVG` did not substantially alter the NMI (slightly increasing from 0.51 to 0.52), the silhouette score range for the dentate gyrus molecular layer (DG ML) is greatly improved after refining the input SVG list with `BatchSVG` (**Figure 1D, Supplementary Figure 5**). Importantly, the DG GCL cluster identified with the `BatchSVG` refined feature set is present in all hippocampal samples and is consistent with the expression of *CALB1* (**Figure 1E**). Finally, we note that common QC metrics do not differ between the DG GCL and ML domains from the original features set versus the refined features set (**Figure 1F**). These results demonstrate the success of `BatchSVG` in reducing the batch-biased impacts and enhancing biologically meaningful spatial domain identification.

### 4.2 Improved spatial domain detection in human dlPFC

This dataset contains human dorsolateral prefrontal cortex (dlPFC) tissue generated using Visium, with a total of twelve samples across three donors [11]. We chose three samples, one from each donor, and identified 1,864 SVGs with `nnSVG`, specifically subsetting to genes that were statistically significant in at least two samples. We used `BatchSVG` with sample as the batch to identify two genes as batch-biased SVGs (**Supplementary Figure 6A-B, Supplementary Table 2**). The `PRECAST` clustering results (*k* = 6 clusters) after using `BatchSVG` are substantially more consistent with known biology than the clusters based on the original feature set (**Supplementary Figures 6C, 7, and 8**). Specifically, the pink and green L3/4/5/6 domains in the pre-`BatchSVG` clustering result separate into two distinct domains expressing L3/4 and L5/6 marker genes, respectively. This improvement is quantified with the increase in NMI comparing either set of domains to manual annotations from 0.39 to 0.58. The silhouette scores for all domains are relatively similar before and after applying `BatchSVG` to the input SVG set (**Supplementary Figure 6D, Supplementary Figure 9**). We again observed that these changes were not associated with differences in QC metrics (**Supplementary Figure 6E**).

## 5 Discussion

We developed `BatchSVG` as a method to identify genes in the SVG set that may be unwanted due to batch effects. We demonstrated that excluding these batch-biased SVGs improves downstream clustering in two SRT datasets and makes the domains more harmonious across samples. Although we have described this method in the context of SRT data, it can also be applied to other contexts. For example, `BatchSVG` can be utilized on a single nucleus RNA-sequencing dataset to identify genes related to the batch variable of sample since the method does not depend on spatial coordinates. The goal of this method is not to replace batch effect correction and integration methods, but to help with interpretability to find genes associated with batch effects in SRT data.

## 6 Supplementary Information

Supplementary materials are available at *Bioinformatics* online.

## 7 Author contributions

**KS:** Conceptualization, Formal analysis, Writing, Visualization; **CH:** Conceptualization, Software, Formal analysis, Writing, Visualization; **JRT:** Conceptualization, Methodology, Formal analysis, Writing, Visualization; **SCH:** Conceptualization, Resources, Writing Review & Editing, Supervision, Project Administration, Funding Acquisition.

## 10 Data and code availability

The open-source software package `BatchSVG` is based on the R programming language. This package is freely available at https://bioconductor.org/packages/devel/bioc/html/BatchSVG.html (at least Bioconductor version 3.21), depending on R (at least version 4.4.0). All data analysis is freely available at https://github.com/kinnaryshah/BatchSVG-analyses.

## References

[1] A. Rao, D. Barkley, G. S. França, and I. Yanai. Exploring tissue architecture using spatial transcriptomics. Nature, 596:211–220, 2021. doi:10.1038/s41586-021-03634-9.

[2] V. Svensson, S. A. Teichmann, and O. Stegle. SpatialDE: identification of spatially variable genes. Nature Methods, 15:343–346, 2018. doi:10.1038/nmeth.4636.

[3] W. Liu, X. Liao, Z. Luo, Y. Yang, M. C. Lau, Y. Jiao, X. Shi, W. Zhai, H. Ji, J. Yeong, and J. Liu. Probabilistic embedding, clustering, and alignment for integrating spatial transcriptomics data with PRECAST. Nature Communications, 14(296), 2023. doi:10.1038/s41467-023-35947-w.

[4] Z. Li, Z. M. Patel, D. Song, G. Yan, J. J. Li, and L. Pinello. Benchmarking computational methods to identify spatially variable genes and peaks. bioRxiv, page 2023.12.02.569717, Dec. 2023. ISSN 2692-8205. doi:10.1101/2023.12.02.569717. URL https://www.ncbi.nlm.nih.gov/pmc/articles/PMC10705556/.

[5] X. Chen, Q. Ran, J. Tang, Z. Chen, S. Huang, X. Shi, and R. Xi. Benchmarking algorithms for spatially variable gene identification in spatial transcriptomics. Bioinformatics, 41(4):btaf131, Mar. 2025. ISSN 1367-4811. doi:10.1093/bioinformatics/btaf131. URL https://academic.oup.com/bioinformatics/article/doi/10.1093/bioinformatics/btaf131/8096371.

[6] G. Yan, S. H. Hua, and J. J. Li. Categorization of 34 computational methods to detect spatially variable genes from spatially resolved transcriptomics data. Nature Communications, 16(1):1141, Jan. 2025. ISSN 2041-1723. doi:10.1038/s41467-025-56080-w. URL https://www.nature.com/articles/s41467-025-56080-w.

[7] F. W. Townes, S. C. Hicks, M. J. Aryee, and R. A. Irizarry. Feature selection and dimension reduction for single-cell RNA-Seq based on a multinomial model. Genome Biology, 21(179), 2020. doi:10.1186/s13059-020-02109-w.

[8] D. Righelli, L. M. Weber, H. L. Crowell, B. Pardo, L. Collado-Torres, S. Ghazanfar, A. T. L. Lun, S. C. Hicks, and D. Risso. SpatialExperiment: infrastructure for spatially-resolved transcriptomics data in R using Bioconductor. Bioinformatics, 38(11):–3, 2022. doi:10.1093/bioinformatics/btac299.

[9] J. R. Thompson, E. D. Nelson, M. Tippani, A. D. Ramnauth, H. R. Divecha, R. A. Miller, N. J. Eagles, E. A. Pattie, S. H. Kwon, S. V. Bach, U. M. Kaipa, J. Yao, C. Hou, J. E. Kleinman, L. Collado-Torres, S. Han, K. R. Maynard, T. M. Hyde, K. Martinowich, S. C. Page, and S. C. Hicks. An integrated singlenucleus and spatial transcriptomics atlas reveals the molecular landscape of the human hippocampus. bioRxiv, 2025. doi:10.1101/2024.04.26.590643.

[10] L. M. Weber, A. Saha, A. Datta, K. D. Hansen, and S. C. Hicks. nnSVG for the scalable identification of spatially variable genes using nearest-neighbor Gaussian processes. Nature Communications, 14 (4059), 2023. doi:10.1038/s41467-023-39748-z.

[11] K. R. Maynard, L. Collado-Torres, L. M. Weber, C. Uytingco, B. K. Barry, S. R. Williams, J. L. C. II, M. N. Tran, Z. Besich, M. Tippani, J. Chew, Y. Yin, J. E. Kleinman, T. M. Hyde, N. Rao, S. C. Hicks, K. Martinowich, and A. E. Jaffe. Transcriptome-scale spatial gene expression in the human dorsolateral prefrontal cortex. Nature Neuroscience, 24:425–436, 2021. doi:10.1038/s41593-020-00787-0.

