## Supplementary Tables and Figures for "BatchSVG: identifying batch-biased genes in the application of spatially variable gene detection"

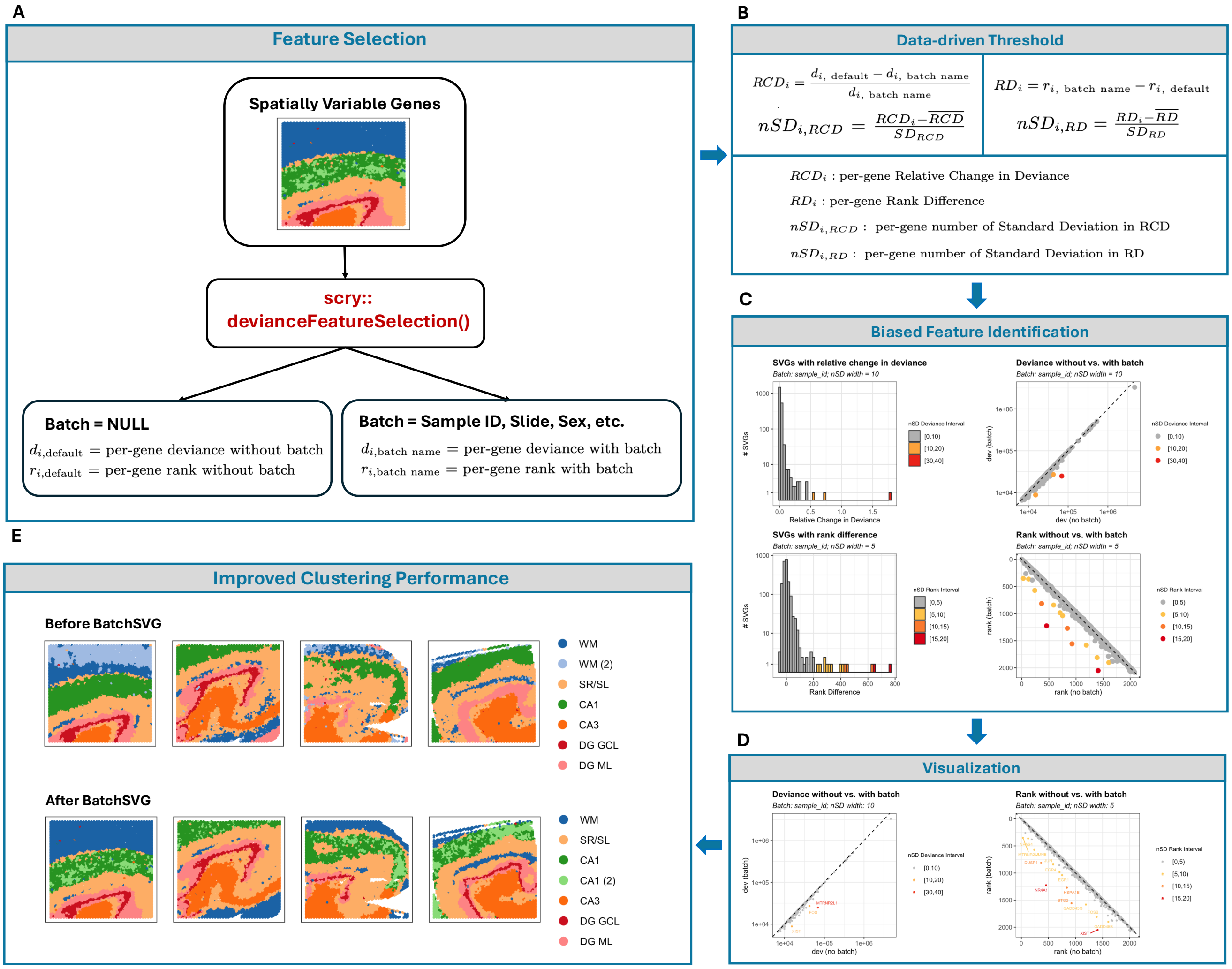


**Figure S1:** The standard workflow of `BatchSVG`: a feature-based quality control method to identify batch-biased spatially variable genes (SVGs). The data is the spatially resolved transcriptomics (SRT) human hippocampus dataset used in the main manuscript. (A) The binomial deviance model is applied to SVGs for per-gene deviance and rank with and without the selected batch variable(s). (B) Data-driven thresholds are computed as the per-gene number of Standard Deviation (nSD) for Relative Change in Deviance (RCD) and Rank Difference (RD). (C) The visualizations help determine the appropriate thresholds for nSD of RCD and RD. (D) `BatchSVG` can identify and visualize batch-biased features based on user-selected thresholds. (E) The refined SVG set, defined by removing batch-biased SVGs, can improve the downstream clustering performance. The clusters were annotated with the following labels - CA: cornu ammonis, DG GCL: dentate gyrus granule cell layer, DG ML: dentate gyrus molecular layer, SR/SL: stratum lucidum and stratum radiatum, WM: white matter.

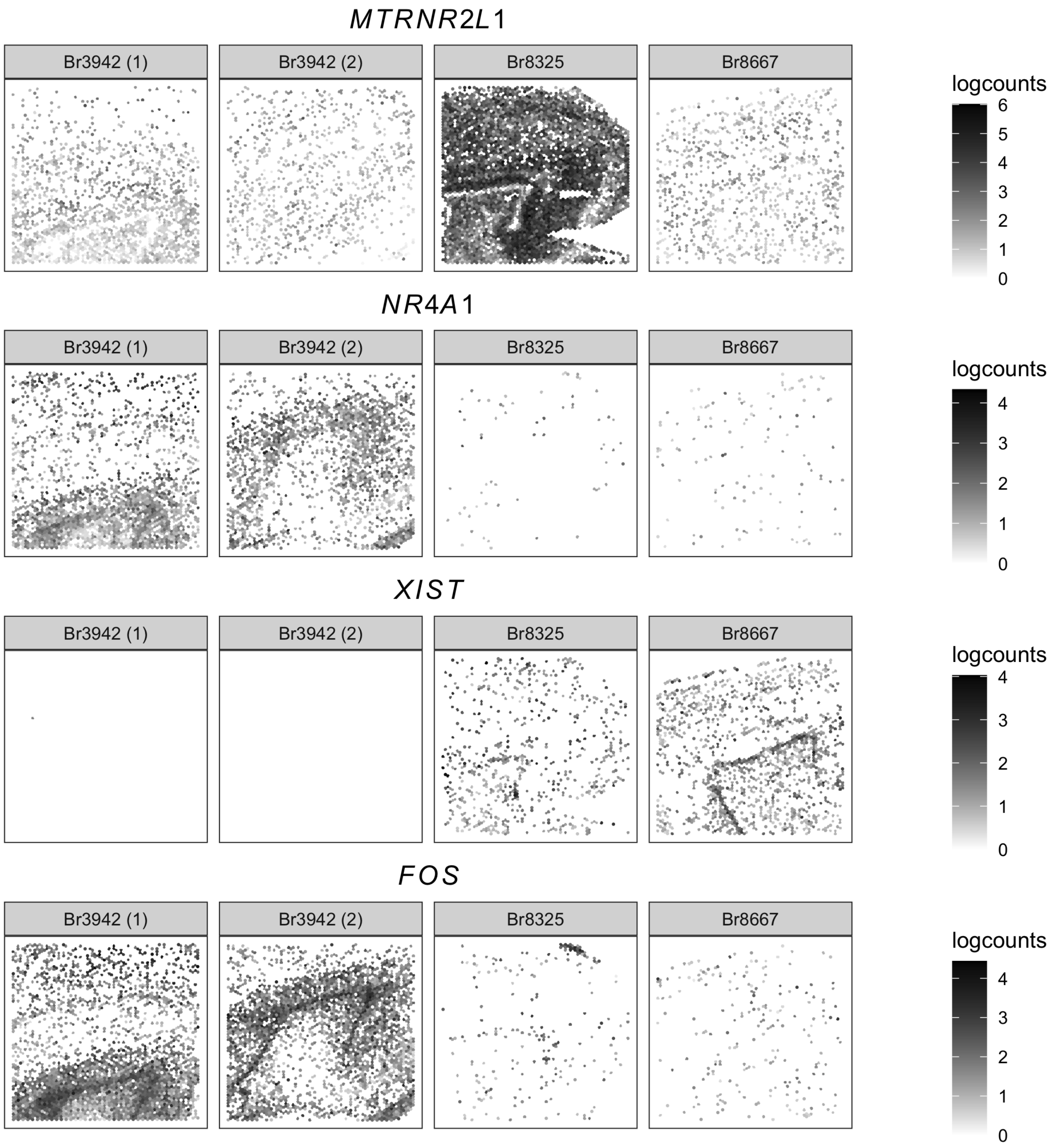


**Figure S2:** Selected spot plots of batch-biased SVGs found in the human hippocampus dataset. The gradient shows the log-normalized counts values for the following genes: *MTRNR2L1*, *NR4A1*, *XIST*, and *FOS*.

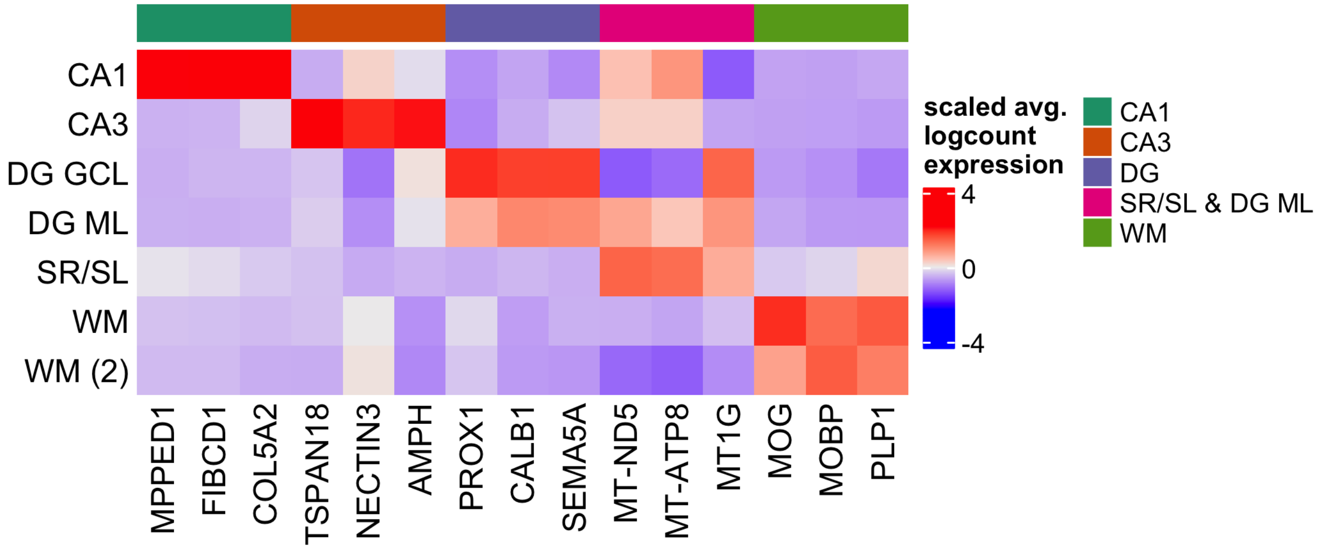


**Figure S3:** Heatmap (centered and scaled) displays hippocampus regional marker genes for domains derived from `PRECAST` when all SVGs were used as input. The marker genes are associated with the following labels - CA: cornu ammonis, DG GCL: dentate gyrus granule cell layer, DG ML: dentate gyrus molecular layer, SR/SL: stratum lucidum and stratum radiatum, WM: white matter.

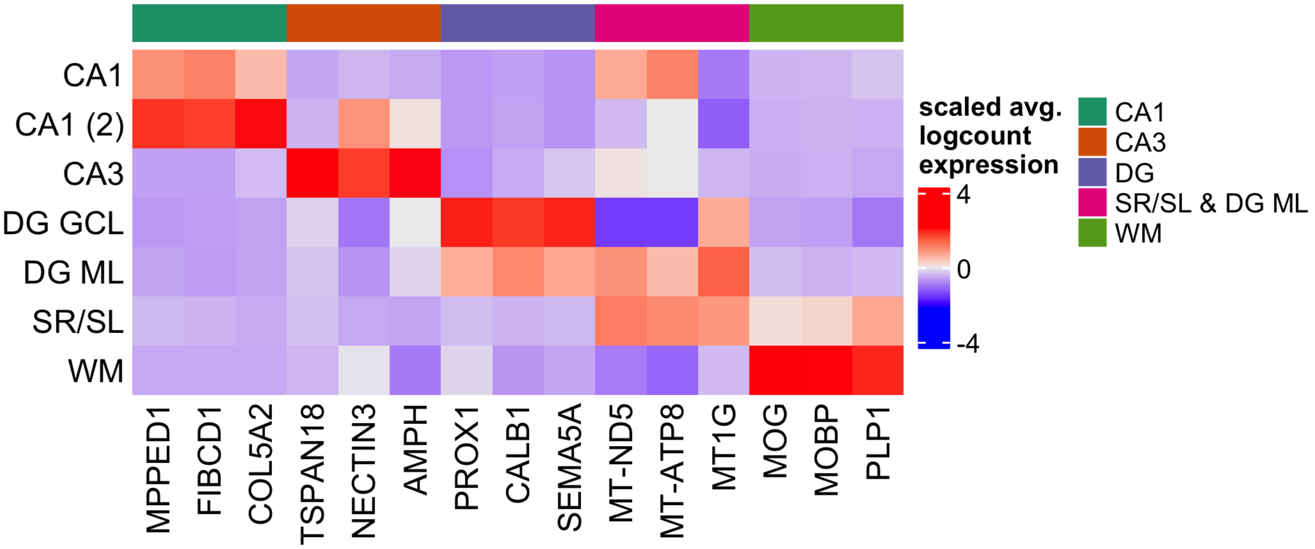


**Figure S4:** Heatmap (centered and scaled) displays hippocampus regional marker genes for domains derived from `PRECAST` when the batch-biased SVGs were excluded from the input SVG list. The marker genes are associated with the following labels - CA: cornu ammonis, DG GCL: dentate gyrus granule cell layer, DG ML: dentate gyrus molecular layer, SR/SL: stratum lucidum and stratum radiatum, WM: white matter.

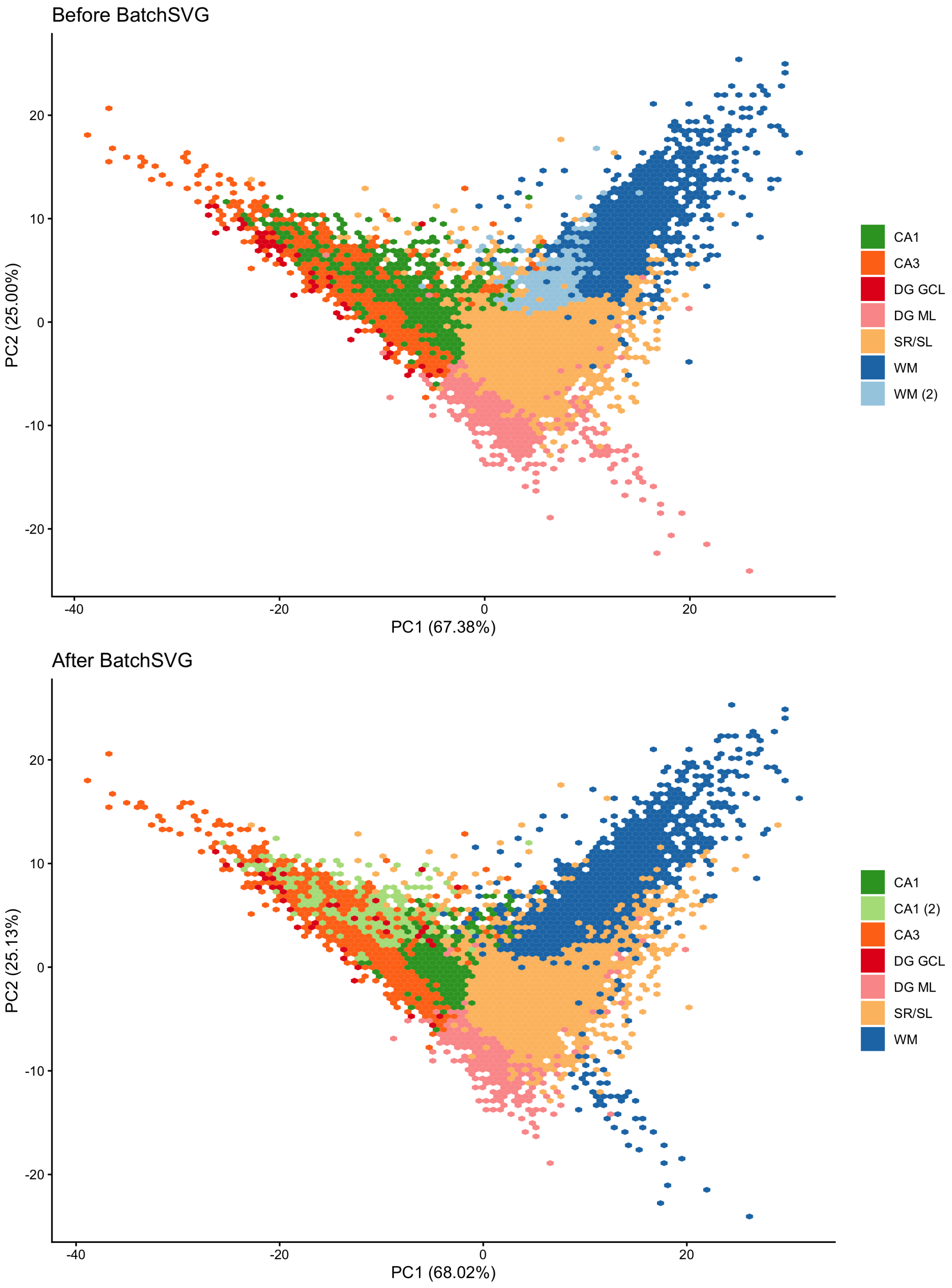


**Figure S5:** Principal components analysis (PCA) plots of the the human hippocampus dataset. The top plot shows the PCA plot of the first two principal components (PCs) for the domains derived from `PRECAST` when all SVGs were used as input. The bottom plot shows the PCA plot of the first two PCs for the domains derived from `PRECAST` when the batch-biased SVGs were excluded from the input SVG list. Color of each hexbin represents the majority cluster. The clusters were annotated with the following labels - CA: cornu ammonis, DG GCL: dentate gyrus granule cell layer, DG ML: dentate gyrus molecular layer, SR/SL: stratum lucidum and stratum radiatum, WM: white matter.

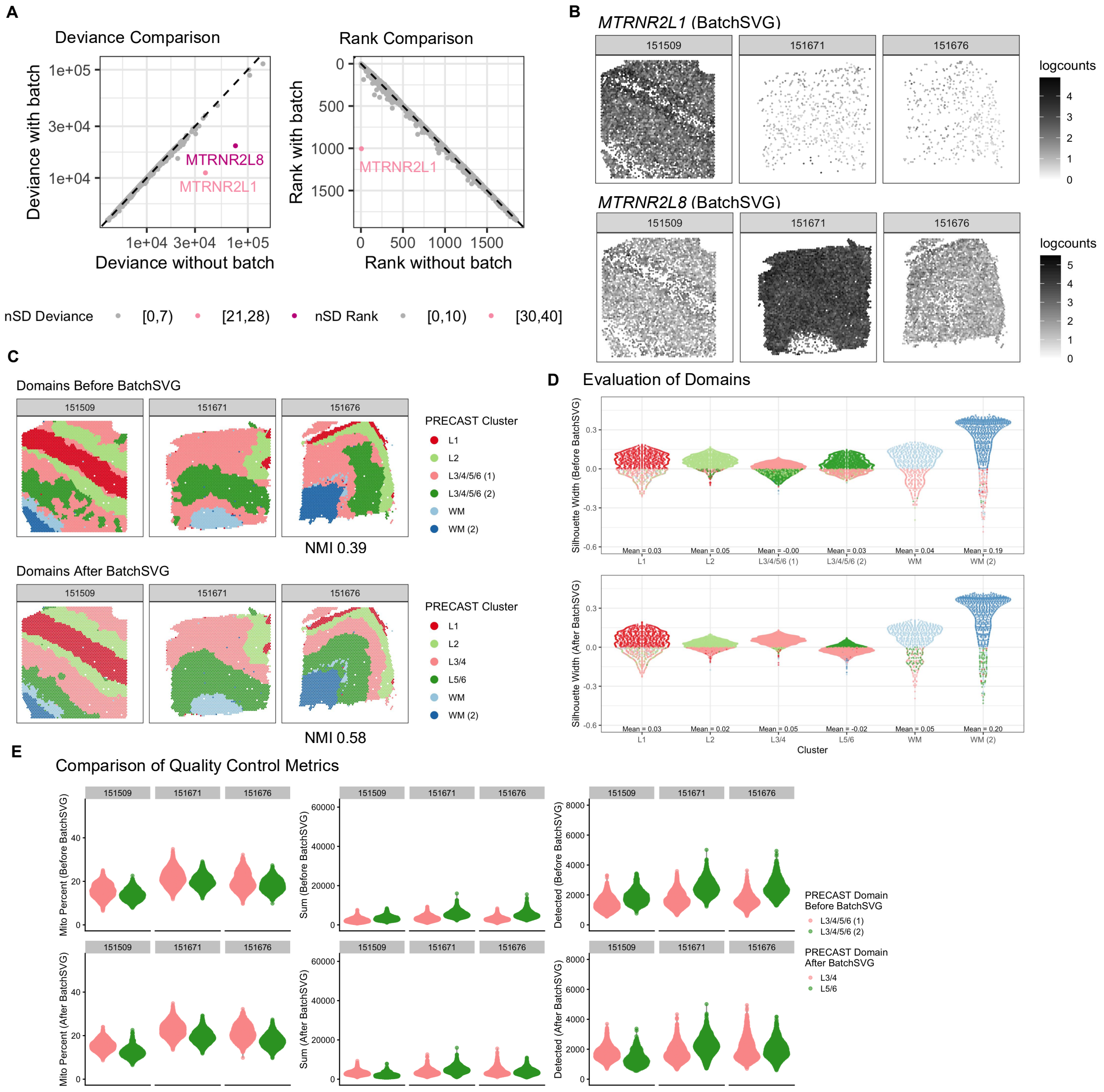


**Figure S6:** Summary figure for dorsolateral prefrontal cortex dataset. (A) Left scatterplot displays the change in deviance for each gene. The $x$-axis is deviance calculated without using any covariates. The $y$-axis is deviance calculated with sample as the covariate. The color represents the number of standard deviations of relative change in deviance. Right scatterplot is similar, substituting rank for deviance. (B) Top spot plot shows the log-normalized counts gene expression for *MTRNR2L1*, which is a batch-biased SVG. Bottom spot plot shows the log-normalized counts gene expression for *MTRNR2L8*, another batch-biased SVG. (C) Top spot plot shows the clusters derived from `PRECAST` when all SVGs were used as input. Bottom spot plot shows the clusters derived from `PRECAST` when the batch-biased SVGs were excluded from the input SVG list. The clusters were annotated with the following labels - L + number: Layer number, WM: white matter. Color represents the cluster. Text displays the normalized mutual information (NMI) for each set of domains compared to manual annotations. (D) Top violin plots display silhouette scores for domains derived from `PRECAST` when all SVGs were used as input. Bottom violin plots display silhouette scores for domains derived from `PRECAST` when the batch-biased SVGs were excluded from the input SVG list. Color represents the closest domain. (E) Violin plots display various quality control metrics for domains derived from `PRECAST` when the batch-biased SVGs were included (top row) or excluded (bottom row) from the input SVG list. Left row displays mitochondrial percent, middle row displays total number of UMIs, and right row displays number of detected genes. Color represents the domain.

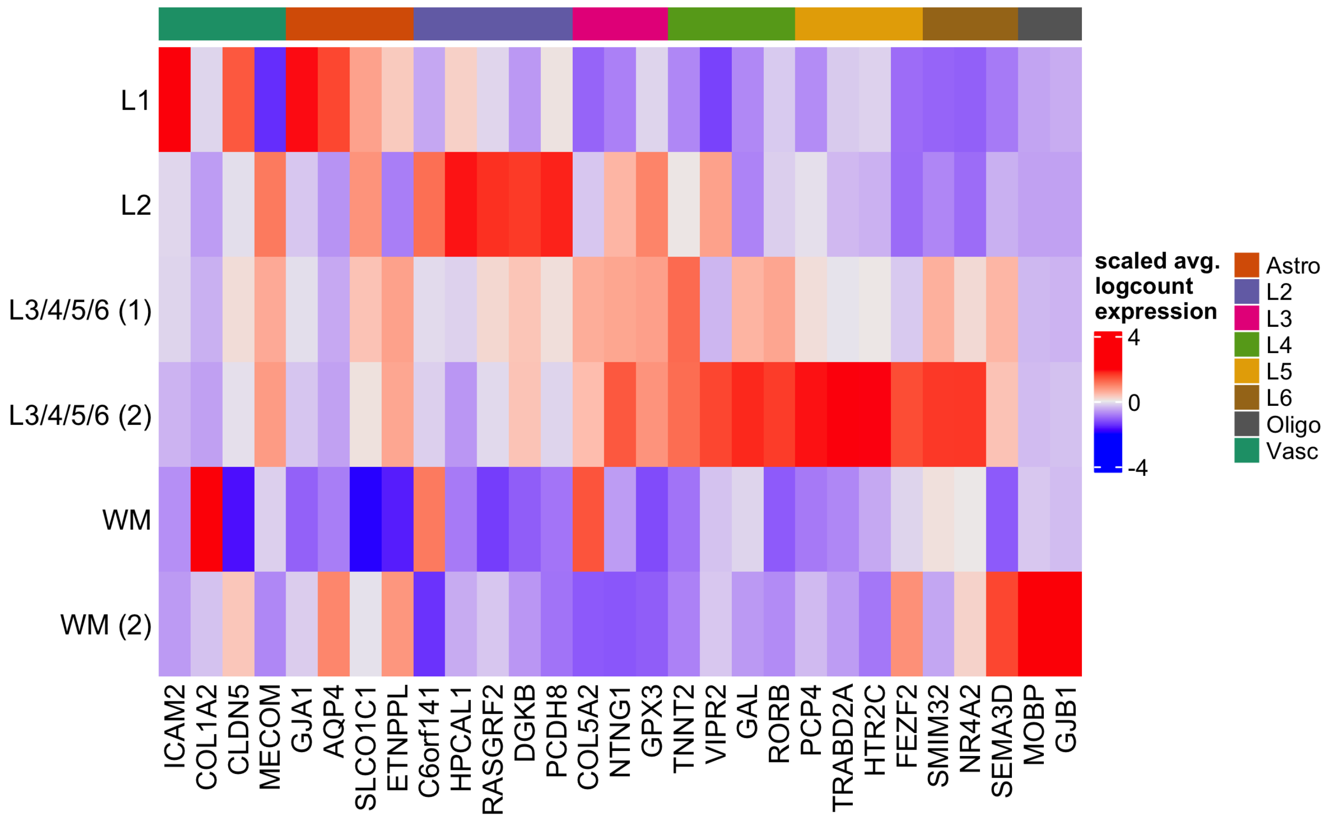


**Figure S7:** Heatmap (centered and scaled) displays dorsolateral prefrontal cortex laminar marker genes for domains derived from `PRECAST` when all SVGs were used as input. The marker genes are associated with the following labels - Astro: astrocyte, L + number: Layer number, Oligo: oligodendrocyte, Vasc: vasculature.

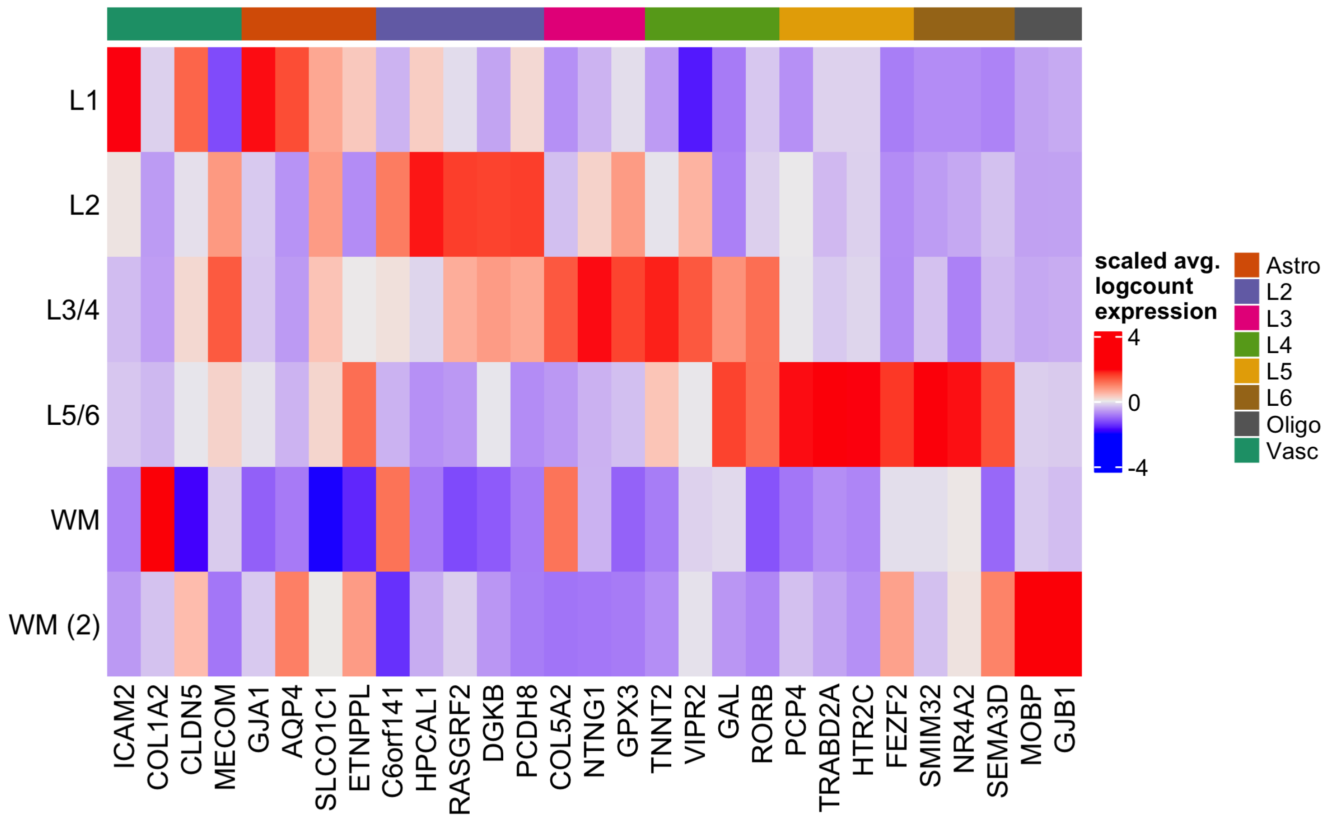


**Figure S8:** Heatmap (centered and scaled) displays dorsolateral prefrontal cortex laminar marker genes for domains derived from `PRECAST` when the batch-biased SVGs were excluded from the input SVG list. The marker genes are associated with the following labels - Astro: astrocyte, L + number: Layer number, Oligo: oligodendrocyte, Vasc: vasculature.

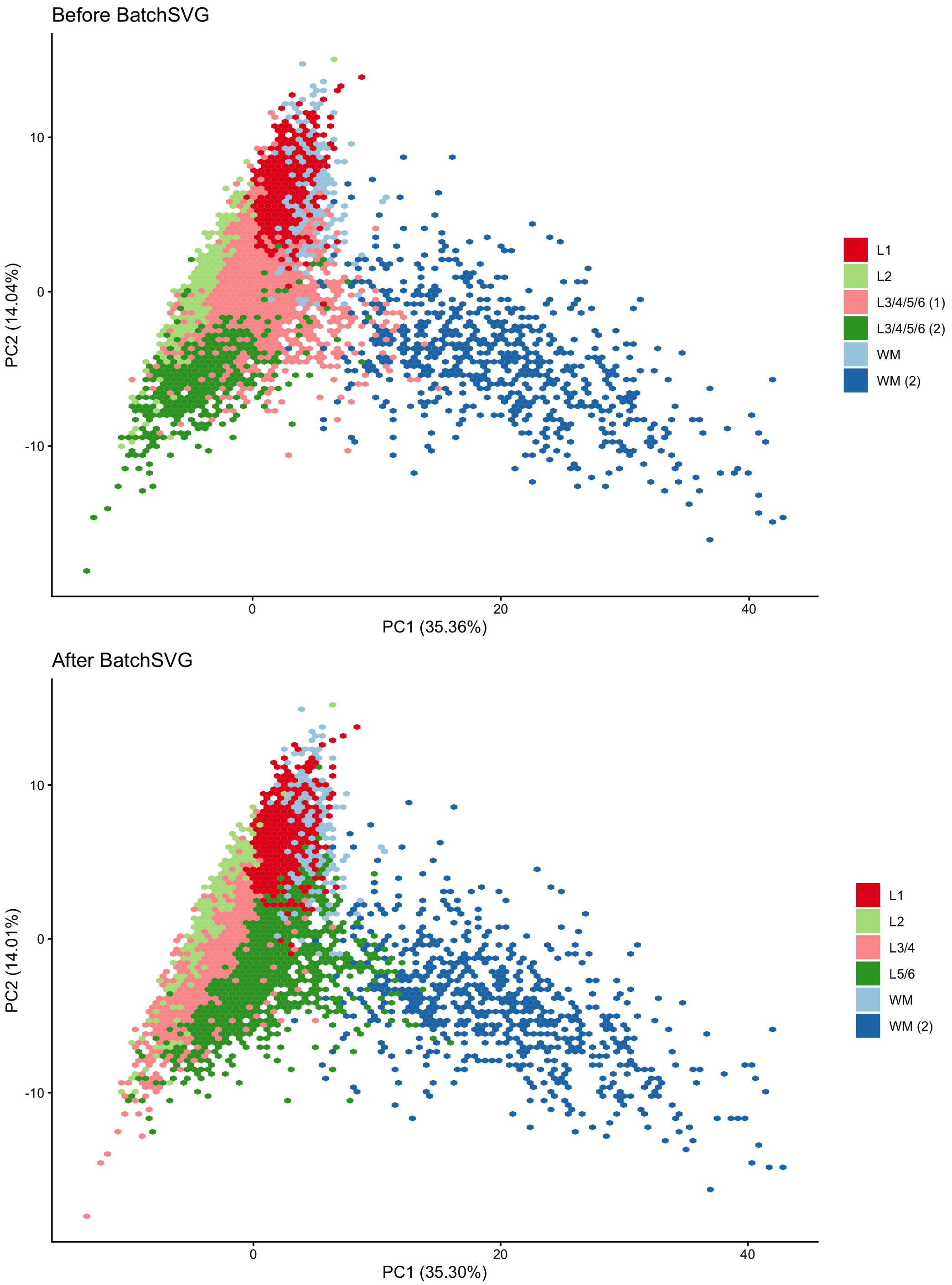


**Figure S9:** Principal components analysis (PCA) plots of the human dorsolateral prefrontal cortex dataset. The top plot shows the PCA plot of the first two principal components (PCs) for the domains derived from `PRECAST` when all SVGs were used as input. The bottom plot shows the PCA plot of the first two PCs for the domains derived from `PRECAST` when the batch-biased SVGs were excluded from the input SVG list. Color of each hexbin represents the majority cluster. The clusters were annotated with the following labels - L + number: Layer number, WM: white matter.

**Table S1:** Human hippocampus batch-biased SVGs. This table shows all genes found to be batch-biased SVGs from the human hippocampus dataset.

| Gene Name | Gene ID |
|---|---|
| JUN | ENSG00000177606 |
| BTG2 | ENSG00000159388 |
| EGR4 | ENSG00000135625 |
| EGR1 | ENSG00000120738 |
| DUSP1 | ENSG00000120129 |
| HSPA1B | ENSG00000204388 |
| GADD45G | ENSG00000130222 |
| NPAS4 | ENSG00000174576 |
| NR4A1 | ENSG00000123358 |
| FOS | ENSG00000170345 |
| MTRNR2L1 | ENSG00000256618 |
| GADD45B | ENSG00000099860 |
| JUNB | ENSG00000171223 |
| FOSB | ENSG00000125740 |
| XIST | ENSG00000229807 |

**Table S2:** Human dorsolateral prefrontal cortex batch-biased SVGs. This table shows all genes found to be batch-biased SVGs from the human dorsolateral prefrontal cortex dataset.

| Gene Name | Gene ID |
|---|---|
| MTRNR2L1 | ENSG00000256618 |
| MTRNR2L8 | ENSG00000255823 |
